# The sheep brain: an interesting translational model for functional MRI studies?

**DOI:** 10.1101/2020.09.02.280024

**Authors:** Nathalie Just, Hans Adriaensen, Pierre-Marie Chevillard, Martine Batailler, Jean-Philippe Dubois, Martine Migaud

**Author notes:** Address : Corresponding author : Nathalie Just, INRAe Centre Val de Loire, UMR Physiologie de la Reproduction et des Comportements, Equipe Néogenèse Hypothalamique, Rôles, Variations et perturbations endocrines - NHyRVana, 37380 Nouzilly, France.

## Abstract

Biomedical investigations in clinically relevant animal models is of crucial interest for faster and appropriate transfer to human. The ovine model has already demonstrated its potential compared to rodents and non-human primates (NHP) in various studies: sheep possess a gyrencephalic brain, captivity is not an issue and can undergo magnetic resonance imaging (MRI) in routine clinical scanners (1.5T, 3T) under identical conditions using similar techniques to humans. To date, the effects of anesthesia have been poorly explored and only few functional MRI (fMRI) studies were conducted in sheep. Here, Blood Oxygen Level dependent fMRI and perfusion MRI were conducted in lambs and adult ewes at 3 T. Robust but weak BOLD responses to visual stimulation were found in the lateral geniculate nucleus (LGN) up to 3% isoflurane anaesthesia. BOLD responses were weaker in adult sheep than in lambs while relative cerebral blood volumes (rCBV) and relative cerebral blood flows (rCBF) were significantly higher in lambs than in adult sheep for both gray and white matter in accordance with previous findings in the human. Assessment of functional responses in healthy individuals under adequate physiological conditions is essential for robust translational studies.

## Introduction

The ovine model has already demonstrated its potential as an effective translational model for biomedical research : several studies were conducted to evaluate various therapies (Schiffner et al., 2018 ; Mitchell et al., 2018 ; Adam et al., 2010) and Magnetic Resonance Imaging (MRI) techniques have been applied successfully in various cases (Stypulkowski et al., 2013, Herrmann et al., 2019). The ovine brain offers several assets:1) sheep is a large animal model with a large brain. These characteristics permit a wide range of procedures that are very difficult in smaller animals among which long-term serial blood or cerebrospinal fluid collections to analyse various parameters in real time over several days (hormonal concentration variation, for example) in conscious and unstressed animals. 2) the ovine brain structure and organization share many similarities with humans: the venous systems are highly comparable and the external cortical surface of the sheep brain is organized into sulci and gyri in contrast to the rodent brain.3) Sheep is a long living species and 4) physiological regulatory processes may share closer kinetics with humans than rodents. In addition, sheep may be easier to handle regarding ethical issues compared to non-human primates (NHP): facilities are widely available across the world including with imaging capacities (Harding, 2017). Moreover, captivity issues as well as contacts with humans are not a problem.

At a time when environmental issues are becoming major areas of investigation, farm animals such as sheep may be of interest for investigating the impact of environmental changes on their nutritional status, their behaviour and their well-being. Notably, sheep are seasonal animals responding to photoperiod. Photoperiodism is a physiological adaptative response to changes in the day length, which is linked in sheep to their sexual activity and reproduction. From late September to early January, short days (shorter duration of luminosity) correspond to a period of intense sexual activity and reproduction while periods from May till July correspond to long days and long duration of luminosity and therefore low sexual activity. Changes occurring during these different periods have been linked to changes in the neurogenic status of several structures of the sheep brain (Batailler et al.,2016; Lévy et al.,2019). However, recent studies report shifts in the duration of long days and short days, which could impact on the sexual behaviour of sheep (Lawrence et al., 2012). In our context, the ovine model showed to be an attractive model for the investigation of adult neurogenesis (Batailler et al., 2016 ; Lévy et al., 2019) and could be an effective way to validate the presence of adult neurogenesis with non-invasive imaging methods such as MR techniques.

In order to study sexual behaviour and hormonal status in the sheep brain, functional magnetic resonance imaging (fMRI) demonstrated valuable features (Ella et al., 2019). Recently, a few studies explored MRI functional possibilities in the sheep brain (Lee et al., 2015; Pieri et al., 2019) demonstrating reproducible and translatable features at clinical magnetic fields (1.5T and 3T). To the best of our knowledge, cerebral perfusion MRI has not been reported in sheep. Lately, there have also been increased interest for a better understanding of the impact of anaesthesia in fMRI preclinical rodent studies (van Alst et al, 2019; Reimann and Niendorf, 2020; Just et al., 2020). Inhaled anaesthetics (isoflurane (ISO), sevoflurane …) in intubated and non-intubated animals remain the most standard type of anaesthesia although many recent reports show dissociation of Blood Oxygen Level dependent (BOLD) and neural responses above 1.2 % ISO anaesthesia (Sonnay et al., 2018; van Alst et al., 2019). The impact of inhaled anaesthetics has been less studied in larger animals. For an appropriate transfer of these studies to the clinical setting where patients are often awake, it is crucial to clearly identify the impact of anaesthetics on the cerebral blood hemodynamic responses. In the present study, we aimed at characterizing BOLD fMRI responses and cerebral perfusion MRI under ISO anaesthesia in the lamb and sheep brains.

## Materials and Methods

### Animals

This study was approved by the Val de Loire animal experimentation ethics committee (CEEAVdL) in accordance with the guidelines of the French Ministry of Agriculture for animal experimentation and European regulations on animal experimentation. All experiments conform with the ARRIVE guidelines. Experiments were performed in accordance with regulations regarding animals (authorization N° 22353 of the French Ministry of Agriculture in accordance with EEC directive). All experiments were performed in 6 adult Ile-de-France (IF) ewes gathered under standard husbandry at the INRA Val-de-Loire research center (Nouzilly, Indre-et-Loire, France, 47°33′00.8”N 0°46′55.3”E). 4 lambs also underwent MRI. The experimental facilities are approved by the local authority (agreement number E37–175–2). Adult ewes were 3 to 4-years old (70-85kg) and the lambs were 4 months old (30-35kg). All animals were fed daily with hay and pellets and had *ad-libitum* access to water.

The day before MRI experiments, each ewe or lamb was transported to the MR facility. Care was taken to always provide company to individual animals to avoid stress since sheep are a gregarious species. Each animal was fasted 24-hours prior to intubation. After immobilization, the sheep was intubated after intravenous administration of a mixture of ketamine and xylazine (20 mg/kg). The jugular vein was catheterized for later injection of 0.1 mg/kg of DOTAREM (Guerbet, Roissy, France) for perfusion MRI imaging. Each ewe or lamb was transported to the MRI room, installed prone on the MRI bed and anaesthesia was immediately switched to 1% ISO in medical air through a respirator (Aestiva, GE Healthcare, Datex-Ohmeda, USA). The respirator allowed continuous control of respiration rates. An oximeter was attached to one of the hind-paws allowing for the control of the partial pressure of oxygen and heart rate. The temperature was controlled through MRI-compatible rectal probe. The duration of each MRI session was 150 minutes for each animal.

#### Magnetic Resonance Imaging

MR imaging was conducted on a 3T whole body MR Scanner (Siemens Verio, Erlangen, Germany) with a large flex coil surrounding the entire head. Following the acquisition of pilot images, T2-weighted multislice images covering the entire brain was conducted for further slice positioning followed by BOLD fMRI.

#### BOLD fMRI

BOLD fMRI was conducted using a multislice single shot gradient echo echo planar imaging (EPI) sequence (TR/TE= 2970/40 ms ; flip angle = 90° ; FOV = 188 ×188 mm^2^ ; Matrix= 72 × 72; Slice thickness = 3 mm ; slices = 20 ; 5 saturation slices were placed around the brain to decrease background signal). A visual stimulus was delivered through a remote -controlled lamp delivering white light and placed at 5 cm distance from the opened right eye (The eyelid was attached using a plastic tool). The paradigm of stimulation consisted of 5 cycles of 30 s OFF-30 s ON periods at a stimulation frequency of 2 Hz for a total acquisition time of 10 minutes. During visual stimulation, the entire MRI room remained in the dark. For each ewe, the isoflurane was increased from 1 to 3 % with a step of 0.5 %. A duration of 5 minutes was allowed between each step.

After BOLD fMRI, structural images covering the entire brain were acquired using the T1-weighted 3 D magnetization prepared rapid gradient echo (MPRAGE) sequence (TR/TE/TI=2500/318/900 ms ; Flip angle = 12; NEX= 2 ; FOV=192 × 192 mm^2^, Matrix size = 384 x384; Voxel size = 0.5 × 0.5 × 0.5 mm^3^).

#### Perfusion

Dynamic susceptibility contrast (DSC) MRI was conducted using a single-shot gradient echo EPI sequence (TR/TE= 1500/21 ms;flip angle = 90° ; FOV : 192 ×192 mm^2^ Matrix size = 64 × 64 ; 19 slices) using a short bolus of 0.1 mmol/kg of DOTAREM (Guerbet, Roissy, France) followed by a saline flush using a power injector (Medrad) injected after 60 s of acquisition at 5ml/s. The acquisition duration lasted a bit more than 2 minutes with 60 to 80 acquisition timepoints.

### Image processing

#### BOLD-fMRI

Functional MR images were processed with SPM12 software (Statistical Parametric Mapping, Wellcome Department of Cognitive Neurology, London, UK) under Matlab R2019b (Matlab, The Mathworks, Natick, MA, USA). Functional MR images were first re-aligned and corrected for motion. Prior to slice-time correction, ewe ‘s images were co-registred to their own structural MPRAGE images and normalized to the in-house developped sheep brain atlas (Ella et al., 2017). Finally, images were spatially smoothed using a 6 × 6 × 3 mm^3^ Gaussian kernel. After these pre-processing steps, the general linear model (GLM) first level analysis was conducted using the canonical hemodynamic response function (HRF) and its time derivative as basis functions. The model included two regressors (the HRF and its temporal derivative) and the motion parameters as nuisance regressors. BOLD responses were mapped as T-value maps overlaid onto anatomical MPRAGE images and onto our-in-house sheep atlas (Ella et al., 2017). Significance of BOLD responses was evaluated at cluster level using FDR -corrected p-value, which gives the family-wise error rate probability due to multiple comparisons and was set to 0.01.

To assess the robustness and reliability of our fMRI results, experiments were repeated 3 weeks after the first session. We also calculated the mean beta values of the BOLD responses in the second session and we qualitatively compared the BOLD response patterns with those obtained in the first session.

#### DSC-MRI

Data were analysed using a Matlab toolbox adapted from Peruzzo et al. (2011) using SVD deconvolution (Zanderigo et al., 2009) and integrating both raw and fitted arterial input functions (AIF).

#### Statistics

A Shapiro-Wilk test was used to test for normality.

Average BOLD responses to each dose of ISO were tested to assess whether they were different from zero using a two-tailed t-test (p<0.05). Differences between BOLD responses, cluster volumes, positions of centers of mass and sessions were assessed using a Kruskal-Wallis test by ranks followed by a Tukey post-hoc test. Differences between average rCBF, CBV, MTT and TTP in gray and white matter between lambs and ewes were also tested with a Kruskal-Wallis test followed by a Tuckey test. The significance level was set to p < 0.05.

## Results

### BOLD fMRI

#### Comparison between lamb and adult ewes responses to visual stimulation

For all animals, visual activation maps were observed within the lateral geniculate nucleus (LGN) contralateral to the stimulated eye. LGN is a structure known as a relay in the thalamus for the visual pathway. It receives sensory inputs from the retina and connects the optic nerve to the occipital nerve in the primary visual cortex (V1). Typical results for one lamb and one ewe are presented in Fig. 1A and 1B respectively. Activated voxels were also seen in cortical areas corresponding to the primary visual cortex (V1) (Fig.1). Single or bilateral negative BOLD activation was also reproducibly found in areas of the brain corresponding to the auditory cortices. Negative BOLD was also found in the Pons (Fig. 1B). For further comparisons, we concentrated on the cluster of activated voxels in LGN.

**Figure 1:**
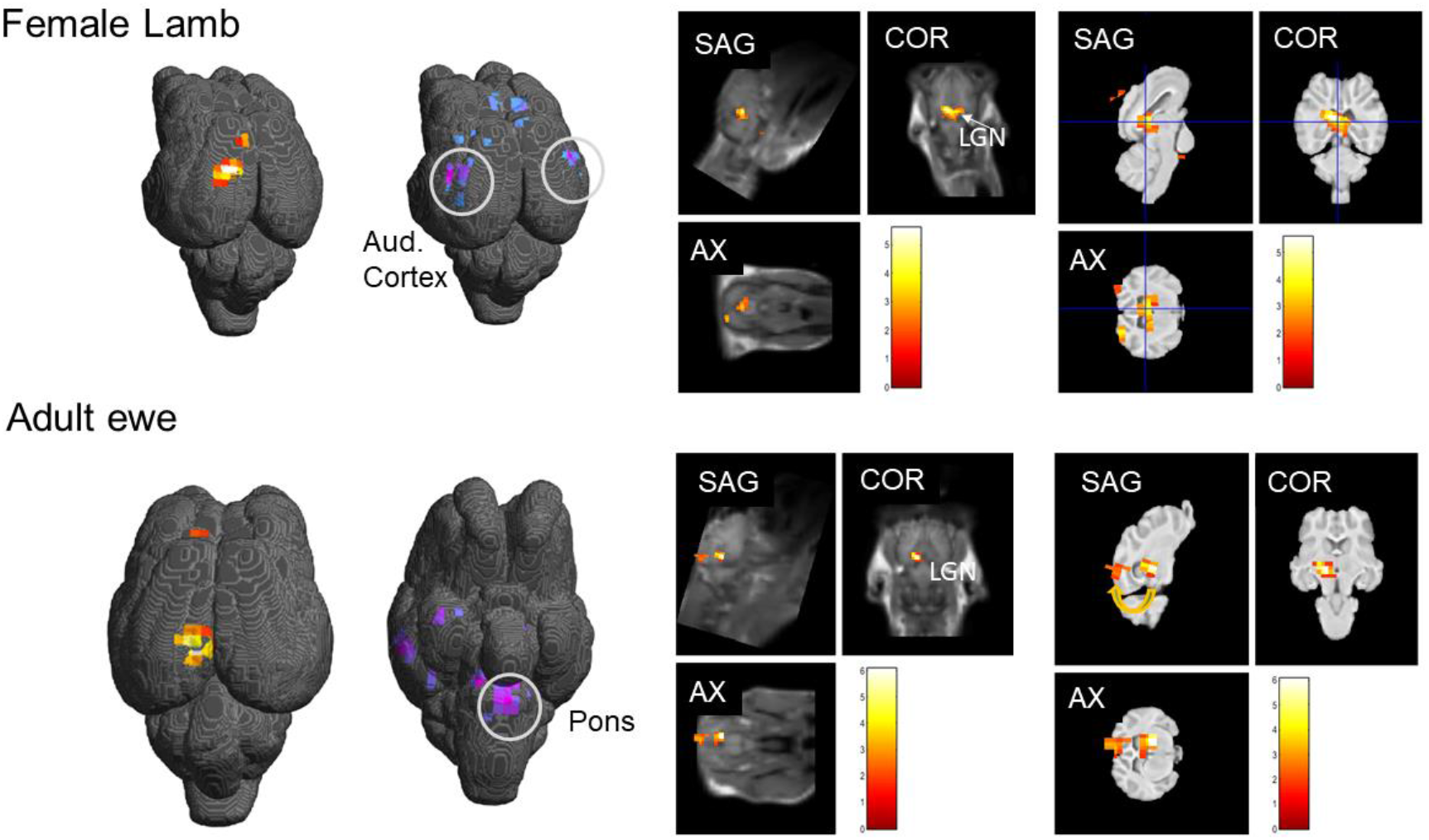
Results of the visual stimulation procedure in the brain of a representative female lamb and representative adult ewe. **A.B.**on inflated brains. Gray circles delineate negative BOLD responses in the auditory cortex and the Pons. For each case BOLD T-value maps were overlaid on **C**.**D**. coregistered sagittal, coronal and axial MPRAGE images and **E**.**F** sheep brain atlas (Ella et al., 2017). The yellow arrow in F. represents the projection of inputs from Lateral Geniculate Nucleus (LGN) to the visual cortex (V1).

#### Impact of the depth of isoflurane anesthesia on BOLD responses in adult ewes

Figure 2 shows T-value maps of activation overlaid onto our in-house sheep brain atlas (Ella et al., 2017) in the axial plane for 1 and 3% ISO in a representative animal. T-value maps demonstrated statistically significant activation in LGN at all levels of ISO. Positive (Fig.2A and 2.C) and negative (Fig.2B and 2.D) BOLD responses are shown. At 3% ISO, the number of positively activated voxels was resduced although this was variable across animals (See Fig.3). Negative BOLD responses were however present in the temporal lobe at all ISO doses. BOLD timecourses within LGN averaged across 6 animals for each ISO dose are depicted in Fig. 3A. No significant difference was found between maximum BOLD percent changes at different doses. Therefore, BOLD timecourses were averaged (Fig.3B) across all ISO doses leading to an average BOLD response of 0.21± 0.08%.

**Figure 2:**
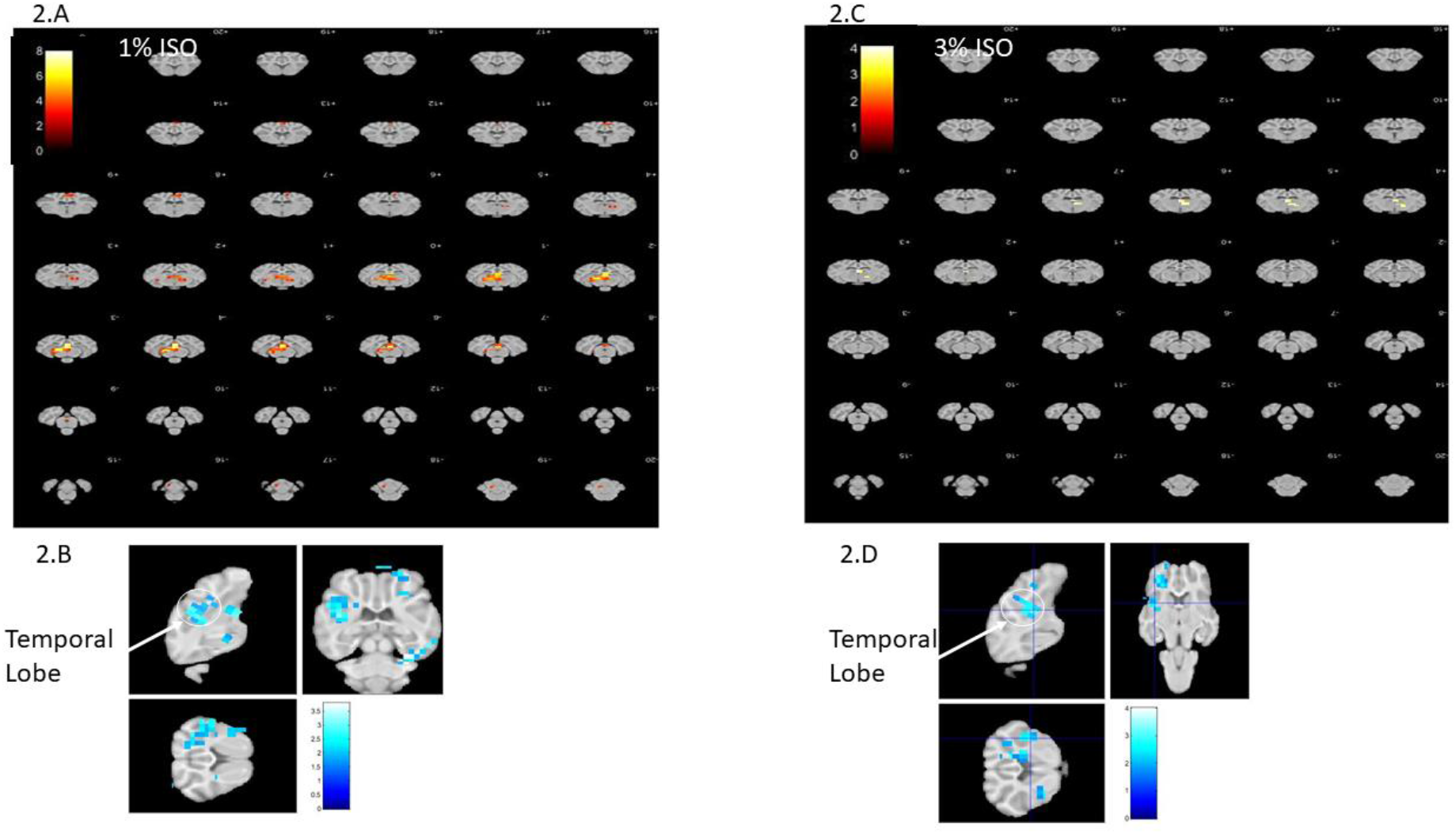
Comparison of BOLD T-value maps overlaid onto our in-house atlas of the adult sheep brain for 1 and 3 % ISO doses within a representative ewe. **A**.Postive BOLD responses in LGN across axial slices at 1% ISO. **B**. Negative BOLD response in the temporal lobe (as indicated) at 1% ISO. **C**. Postive BOLD responses in LGN across axial slices at 3% ISO. The number of activated voxels was significantly reduced in this animal at 3% ISO. **D**. Negative BOLD response in the temporal lobe (as indicated) were still present at 3% ISO

**Figure 3:**
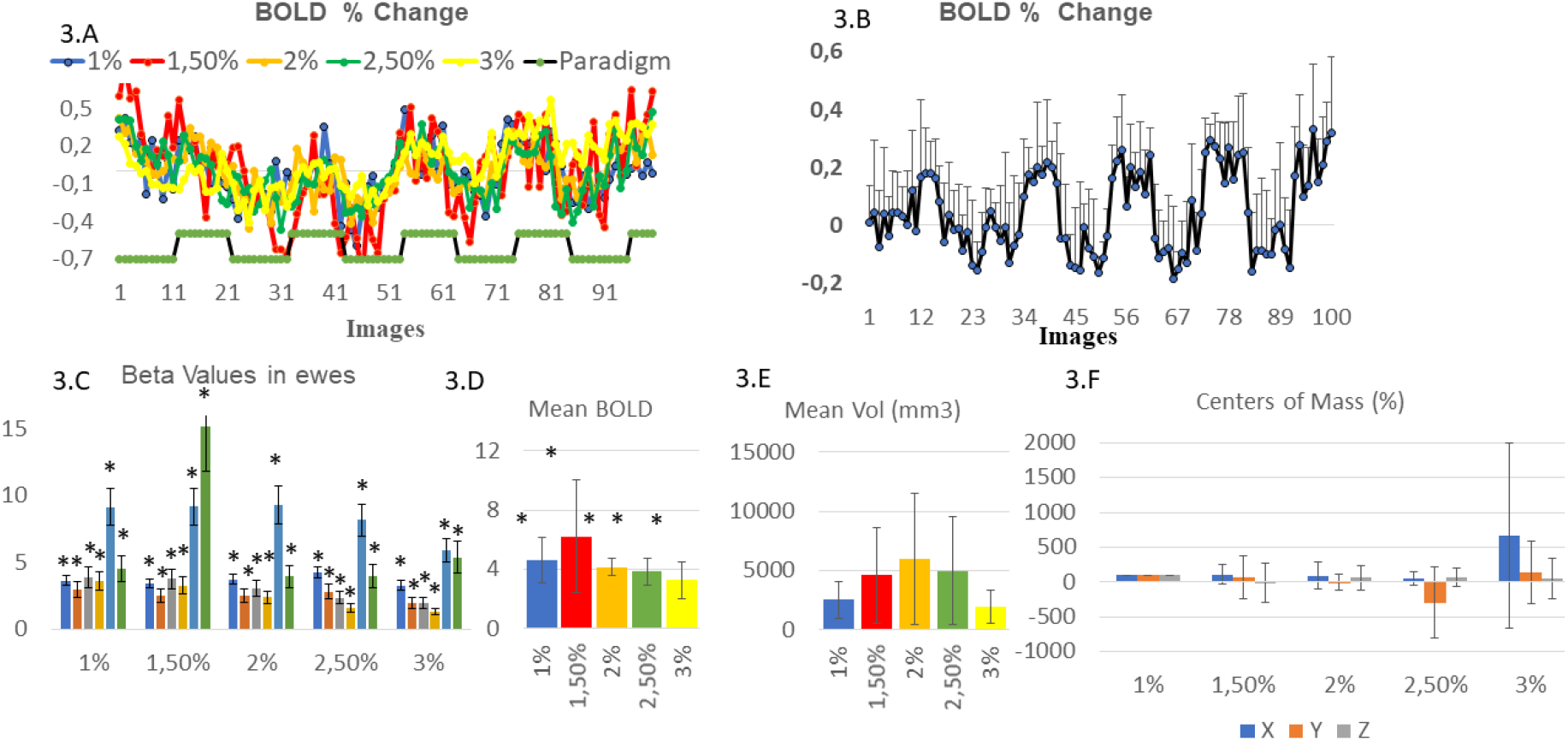
Influence of ISO dose on BOLD responses. **A.** BOLD timecourses at increasing ISO doses and paradigm of visual stimulation. No significant difference was detected. **B**. Average BOLD timecourse (± standard deviation) showing robust but low BOLD responses in LGN.**C**. Average BOLD response (β values) to visual stimulation in each adult sheep as a function of ISO dose. Asterisks indicate values significantly different from zero (p<0.05). Error bars show the standard error. **D**. Mean BOLD responses (averaged across animals). Asterisks indicate values significantly different from zero (p<0.05). Error bars show the standard error. **E**. Mean cluster volumes as a function of ISO dose. **F. Mean change of the coordinates of the centers of mass as a function of ISO dose**. At 1% ISO coordinates X, Y and Z in our sheep atlas space were set to 100% and relative changes were calculated for increasing ISO doses.

To investigate the impact of the ISO dose on the BOLD response to visual stimulation, the mean BOLD response for each ISO dose (represented by the β values for the relevant GLM regressor) was computed. Results (see Fig. 3C) showed a trend towards a reduction of β value amplitudes above 1% ISO except for 2 animals showing maximal β values at 1.5% ISO. On average, no significant differences were found beween BOLD responses between 1 and 3 % ISO (Fig. 3D). Again, individual LGN cluster volumes demonstrated significant dispersion across animals and no significant difference between mean cluster volumes (p= 0.3) (Fig. 3E). However, single animals presented a trend towards increased volumes at 1.5% and 2% ISO followed by a decrease up to 3 % ISO.

#### Changes in the positions of the centers of mass as a function of isoflurane dose

To further evaluate the impact of ISO on BOLD responses, the mean coordinates of the centers of mass were evaluated for each animal and extracted using SPM12 functionalities. On average, no significant differences of the positions of the centers of mass as the dose of ISO increased were found compared to the lowest dose of 1% owing to large variabilities across animals (Fig.3F) although important changes can be observed for X (+ 666 %) and Y (−300% and +141%) centers of mass at 2.5 and 3% ISO and significantly increased standard deviations at these ISO doses.

Positions of centers of mass of LGN activation were also evaluated from images acquired at 3 weeks difference in each animal demonstrating significant deviations (p<0.001) at all ISO doses compared to 1% ISO independent of the scanning timepoint (Fig.5A and 5B). This result suggested an impact of ISO anaesthesia on the position of activation clusters with random effects on their orientation.

**Figure 4:**
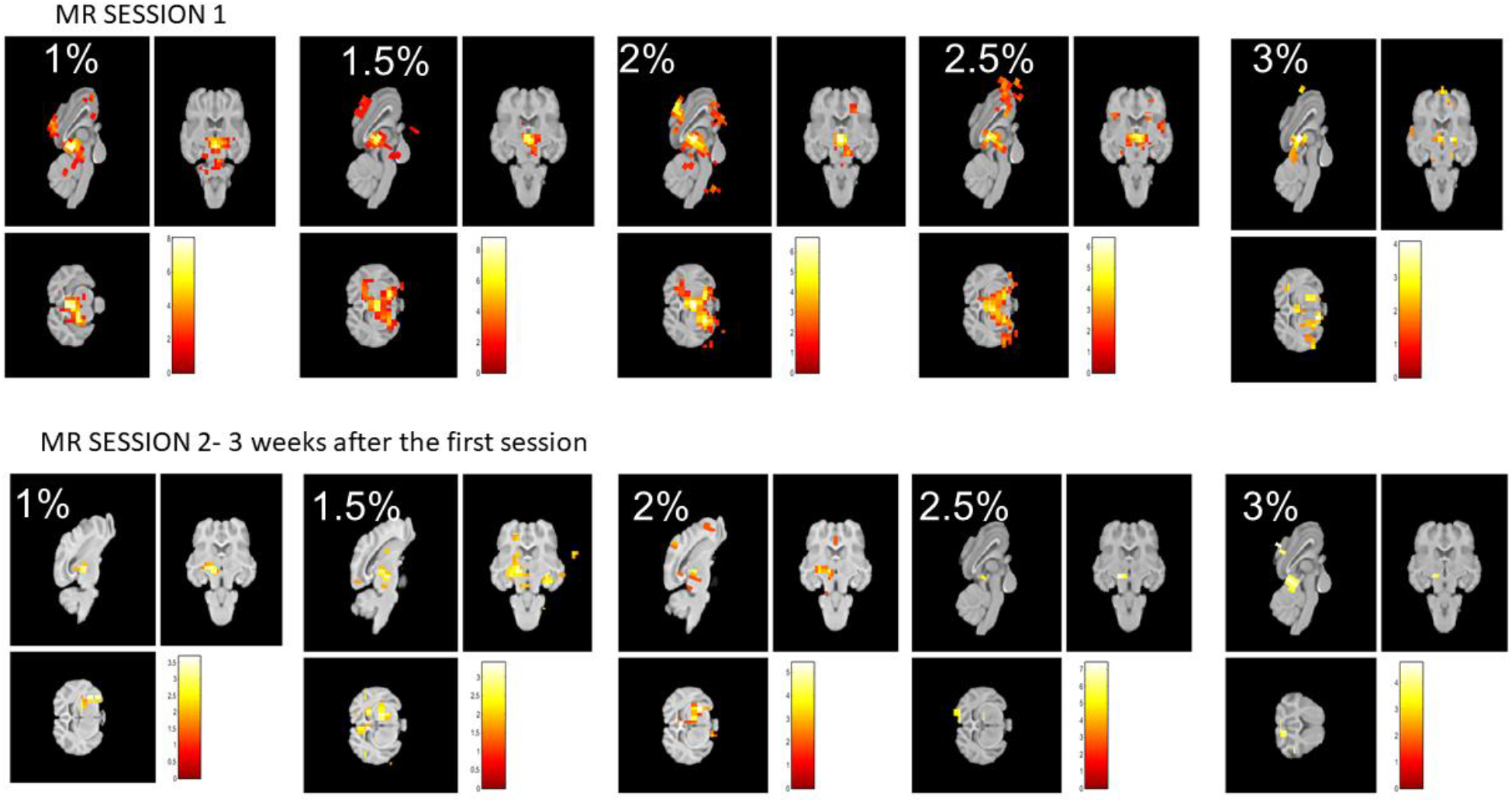
Comparison of BOLD T-value maps acquired at 3 weeks interval for increasing doses of Isoflurane from 1% to 3% within a representative animal.

**Figure 5:**
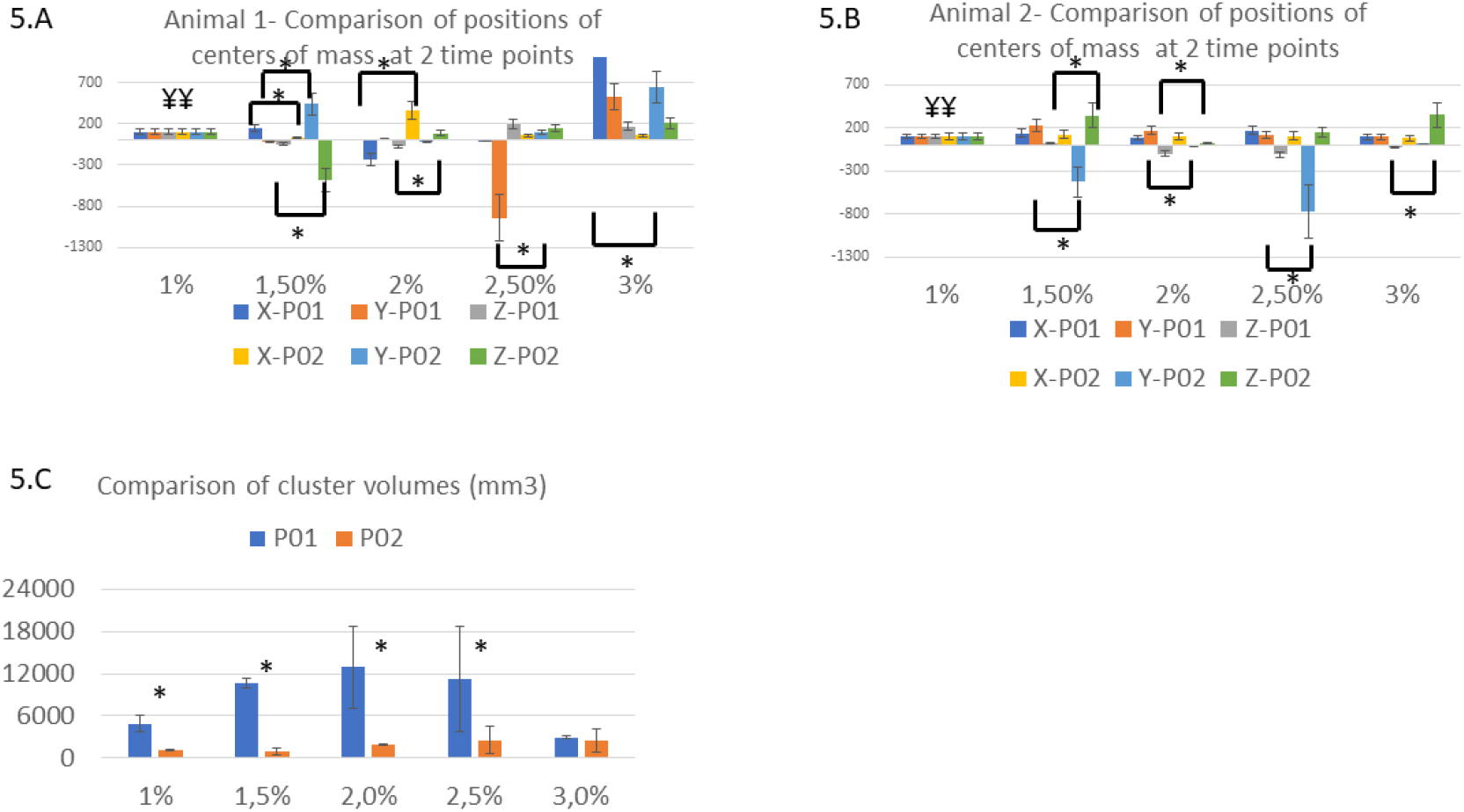
A. B. Comparison of position of centers of mass and cluster volumes in animals scanned at 3 weeks interval. As in Fig.3, coordinates X, Y and Z were set to 100% at 1% ISO and relative changes were calculated. ¥¥ represents significant differences (p<0.05) of coordinates at 1% relative to all other doses. * represents significant differences between first and second MR session coordinates. **C. Comparison of cluster volumes between MR sessions**. * p<0.05

Cluster volumes were significantly reduced between the first and second scanning session up to 2.5 % ISO (Fig.5C).

#### Comparison between lambs and ewes

Fig. 6A compares average β values between lambs for a 1 % Isoflurane anesthesia (ISO) demonstrating no significant changes owing to the large distribution of values across lambs (Fig. 6 A). Mean β values were higher in lambs than in ewes (13.6 ± 8.5 versus 4.6 ± 1.5 ; p=0.08) (Fig. 6 B). Mean volumes of the LGN activated clusters (± standard deviations) were also not significantly different between lambs and ewes (p = 0.8) (Fig. 6 B).

**Figure 6:**
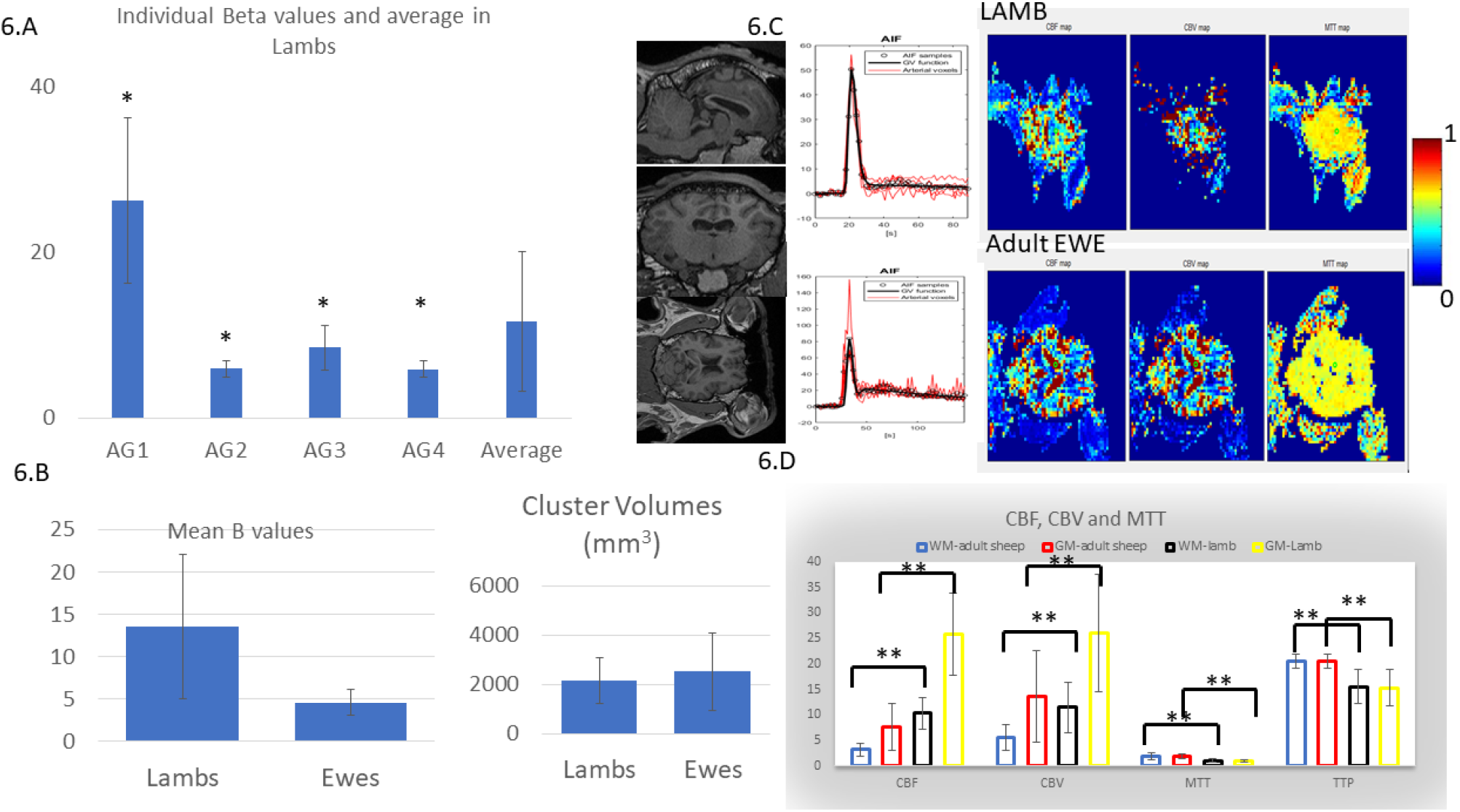
Comparison of BOLD responses and Perfusion MRI in lambs and ewes. **A**. Average BOLD responses (β values) in lambs and average across animals (n=4). Asterisks indicate values significantly different from zero (p<0.05). Error bars show the standard error. **B. Comparison of BOLD responses and cluster volumes between lambs and ewes**. No significant differences were found between mean beta values (p=0.08) and cluster volumes (p= 0.8). **C**. Examples of relative perfusion maps (CBF, CBV and MTT maps) in one lamb and one ewe in the coronal plane. Anatomical MPRAGE images are shon for reference as well as AIFs automatically detected and fitted for deconvolution **D**. Comparison of rCBF, rCBV, MTT and TTP in white matter and gray matter between lambs (n=4) and ewes (n=6). ** p<0.001. Standard errors are represented.

### Perfusion MRI

Finally, perfusion acquisitions were performed both in lambs and adult ewes in order to assess resting-state perfusion parameters : relative cerebral blood flow (rCBF), relative cerebral blood volume (rCBV) and mean transit times (MTT). Fig. 6C shows voxel by voxel rCBF, rCBV and MTT maps obtained in lambs and ewes (Coronal views). Regions of interest (ROIs) encompassing 5 pixels were drawn manually across the whole brain in white matter (WM) and gray matter (GM) in 4 lambs and 4 ewes. Fig. 6D compares mean perfusion parameters between WM and GM. rCBF and rCBV were significantly higher in lambs than in ewes for both WMand GM (p< 0.0001) while MTT and the time to peak (TTP) were significantly smaller in lambs than in ewes (p<0.0001). Mean rCBF and mean rCBV values in GM were 2.5 times the values of WM in both lambs and ewes.

## Discussion

The ovine model represents a very attractive alternative to rodents and NHP for translational neuroimaging studies with its gyrencephalic brain presenting similar vascular features to the human brain. Moreover, ethical concerns are less problematic than for NHP. Therefore, the ovine model could develop into an extremely relevant model for investigating the environmental impact on brain functions with neuroimaging techniques.

All these features comfort us in our idea to show that sheep is a clinically relevant model, which can be studied *in-vivo* in clinical MRI scanners under very similar conditions to the ones experienced by humans.

### Impact of Isoflurane anesthesia on BOLD fMRI responses in sheep

In the present work, BOLD fMRI and perfusion studies were conducted in 3 months and half old lambs and in adult ewes in order to evaluate the feasibility and reproducibility of BOLD functional brain activation in the sheep under isoflurane anesthesia. Robust and repeated activation of LGN was found across animals, across sessions and up to 2 % ISO. However, cortical activation was not systematically observed. The function of LGN during visual processing has been extensively studied both in animals and in human studies (Chen et al., 1998). LGN is the primary target of retinal afferents crossing the optic chiasm. The LGN itself then projects to the visual cortex V1. LGN represents a suitable target to explore activation of deeper structures such as the thalamus. In the present work, robust and repeated activation of LGN was found across animals and across sessions. Currently, mean BOLD percent changes were weak (0.21± 0.08% with maximum BOLD responses of 0.5% for an ISO dose of 1.5%) in accordance with a previous study in the human LGN (Chen et al., 1998). In our context, weak BOLD reponses in LGN may arise from the use of ISO (discussed below) and/or the limited range of frequencies of stimulation delivered during our visual paradigm. Current developments are under way in our institution to deliver adapted visual stimuli with dedicated instrumentation (Sonnay et al., 2018). A previous study in sheep (Lee et al., 2015) reported robust visual activation in the primary visual cortex (V1) with BOLD responses around 2%. However, in this previous study, sheep were anesthetized with N-methyl-D-aspartate (NMDA) receptor antagonist, Telazol, in order to precisely avoid alteration of cerebral blood hemodynamic responses due to volatile anesthesia such as ISO.

In contrast to humans, anesthesia remains a pre-requesite for preclinical MRI investigations. Inhaled gas remains one of the standard anesthesia in preclinical MRI studies for its ease of use (intubation is not a necessity and no canulation is required) and fexibility in dosage. However, in rodent BOLD fMRI studies, ISO anesthesia demonstrated many disadvantages compared to other anesthetics (i.e medetomidine) as BOLD responses disappear due to inherent vasodilatatory effects and increasing CO2 levels in non-ventilated animals (Reimann H et al., 2020 ; van Alst et al., 2019). In sheep, the effects of anesthesia for preclinical MRI studies has not been investigated to the best of our knowledge. With the current setup for visual stimulation, BOLD amplitudes in LGN of intubated sheep demonstrated no significant differences from 1% to 3% ISO. In terms of β values, no significant difference was measured as a function of ISO dose. Comparison of cluster volumes across doses did not reveal significant differences either. On the other hand, significant differences were revealed between MR sessions showing a significant decrease of cluster volumes at all doses several weeks after the first MR session. Adaption of neuronal populations may explain this phenomenon and is not surprising given the retinotopic organization of the LGN (Chen et al., 1999) consisting of several layers of magnocellular and parvocellular cells. Under anesthesia, reduced afferent input and the enhanced contribution of inhibitory systems reduce the spread of stimulus-related activity, resulting in more focal activation and correspondingly smaller BOLD response area (Aksenov et al., 2015). Moreover, exposure to ISO was reported to induce randomized, uncoordinated neuronal activity within an inherently well-organized circuit. In addition, the position of centers of mass demonstrated increased deviations as the dose of ISO increased. These deviations were more prominent at 2.5 and 3 % ISO but large variabilities were also detected within the adult sheep population already at 1.5% ISO compared to the low dose of 1% ISO. The influence of ISO on the position of the center of mass was further increased on subsequent fMRI sessions with increased cluster shifts seen already at 1.5% ISO. Exploration of literature did not reveal similar investigations in the human or rodent brain. However, the reliability of the center of mass parameter for the accurate localization of functional responses was investigated revealing the necessity of high threshold levels for an increased reliability (Fesl et al., 2008 ; Swallow et al., 2003). In addition, functional localization was more reliable when performed on each individual using data in atlas space rather than grouping. Our results also suggest lower functional localization reliability at high ISO doses (2.5 and 3% ISO) (see Fig. 2 and 4), which is why we avoided grouping analysis.

### Comparison of BOLD and perfusion responses between lambs and adult ewes

For a better understanding of developmental and aging brain processes and their alterations as well as an improved assessment of the origins of abnormalities of cognitive and psychiatric brain functions, comparisons of functional neuroimaging evaluations across ages is more and more necessary (Ghosh et al., 2010 ; Mc Gregor et al., 2012 ; Restom et al., 2007). In this regard, preclinical imaging studies in sheep may be more informative than in rodents owing to relatively long cerebral development from childhood till adulhood and longer life expectancy. Moreover, the ovine model has shown interesting features for the evaluation of neurodevelopmental disorders (Wahl, 2004 ; Ramadoss et al., 2008) and there is increasing interest in their behaviour including learning, cognition, emotions and social complexity (Marino and Merskin, 2019). In this context, BOLD responses to visual stimulations were compared in the LGN of lambs and ewes revealing a 3-fold BOLD response (β value) in lambs compared to ewes. Cluster activation sizes were not different in lambs and ewes (not normalized to the whole brain volumes) at an ISO dose of 1.5%. For practical and ethical reasons, notably increased death rate at higher doses, the influence of ISO at different doses was not investigated in lambs.

Finally, MR perfusion studies were conducted successfully in both lambs and ewes. Assessment of relative CBV, CBF and MTT in gray and white matter is reported here for the first time to the best of our knowledge. A 3.5 fold lower rCBF and a 2-fold lower rCBV in both GM and WM were found on average in adult ewes compared to lambs. MTT values were lower in lambs than ewes. These findings are in agreement with findings in the human brain (Wu et al., 2016). Assessment of baseline cerebrovascular values are of interest for future studies of brain lesions (stroke, cancer) and in ovine models of diseases allowing an improved interpretation of BOLD responses.

## Conclusion

Here, BOLD fMRI and MRI perfusion findings are reported in sheep. The ovine model represents an interesting translational model, which can be investigated under identical conditions to humans in routine clinical MR scanners at a field strength of 3T. Similar cerebral features to human were widely described in sheep, in particular their gyrencephalic brain making of them an effective translational model compared to rodents and an appropriate alternative to NHP.

## Author contributions

NJ and MM designed the study. NJ and HA conducted the MR experiments. NJ analyzed and interpreted the data. MB and JPD contributed to animal experiments. NJ wrote the manuscript. All authors read, corrected and approved the manuscript.

## Acknowledgments

Authors would like to thank Dr Arsène Ella and Dr Matthieu Keller for providing their atlas of the sheep brain. This study was funded by a grant from Agence Nationale de la Recherche (ANR-16-CE37-0006-01) to Martine Migaud.

## References

Adam CL, Findlay PA Decreased blood-brain leptin transfer in an ovine model of obesity and weight loss: resolving the cause of leptin resistance. Int J Obes (Lond). 2010 Jun;34(6):980–8. doi: 10.1038/ijo.2010.28.

Aksenov DP, Li L, Miller MJ, Iordanescu G, Wyrwicz AM. Effects of anesthesia on BOLD signal and neuronal activity in the somatosensory cortex. J Cereb Blood Flow Metab. 2015 Nov;35(11):1819–26. doi: 10.1038/jcbfm.2015.130.

Batailler B, Derouet L, Butruille L, Migaud M. Differential effects of oxytocin on olfactory, hippocampal and hypothalamic neurogenesis in adult sheep.Brain Structure and Function 221 (6), 3301–3314

Chen W, Kato T, Zhu XH, Strupp J, Ogawa S, Ugurbil K. Magn Reson Med. Mapping of lateral geniculate nucleus activation during visual stimulation in human brain using fMRI. 1998 Jan;39(1):89–96. doi: 10.1002/mrm.1910390115.

Chen W, Zhu XH, Thulborn KR, Ugurbil K. Retinotopic mapping of lateral geniculate nucleus in humans using functional magnetic resonance imaging. Proc Natl Acad Sci U S A. 1999 Mar 2;96(5):2430–4. doi: 10.1073/pnas.96.5.2430.

Ella A, Delgadillo JA, Chemineau P, Keller M. Computation of a high-resolution MRI 3D stereotaxic atlas of the sheep brain. Journal of Comparative Neurology. 2017. 525 (3), 676–692

Ella A, Barrière DA, Adriaensen H, Palmer DN, Melzer TR, Mitchell NL, Keller M. The development of brain magnetic resonance approaches in large animal models for preclinical research. Anim Front. 2019 Jun 25;9(3):44–51. doi: 10.1093/af/vfz024. eCollection 2019 Jul.

Fesl G, Braun B, Rau S, Wiesmann M, Ruge M, Bruhns P, Linn J, Stephan T, Ilmberger J, Tonn JC, Brückmann H Is the center of mass (COM) a reliable parameter for the localization of brain function in fMRI? Eur Radiol 2008;18(5):1031–7. doi: 10.1007/s00330-008-0850-z.

Ghosh SS, Kakunoori S, Augustinack J, Nieto-Castanon A, Kovelman I, Gaab N, Christodoulou JA, Triantafyllou C, Gabrieli JD, Fischl B. Evaluating the validity of volume-based and surface-based brain image registration for developmental cognitive neuroscience studies in children 4 to 11 years of age. Neuroimage. 2010 Oct 15;53(1):85–93. doi: 10.1016/j.neuroimage.2010.05.075.

Harding JD. Nonhuman primates and translational research: progress, opportunities and challenges. ILAR J. 2017;58(2),141–150.

Herrmann AM, Cattaneo GFM, Eiden SA, Wieser M, Kellner E, Maurer C, Haberstroh J, Mülling C, Niesen WD, Urbach H, Boltze J, Meckel S, Shah MJ Development of a Routinely Applicable Imaging Protocol for Fast and Precise Middle Cerebral Artery Occlusion Assessment and Perfusion Deficit Measure in an Ovine Stroke Model: A Case Study.. Front Neurol. 2019 Nov 14;10:1113. doi: 10.3389/fneur.2019.011

Just N. Proton functional magnetic resonance spectroscopy in rodents. NMR Biomed. 2020 Jan 22:e4254. doi: 10.1002/nbm.4254.

Lawrence TLJ, Fowler VR, Novakofski JE. Growth of farm animals 3rd Edition Lee W, Lee SD, Park MY, Foley L, Purcell-Estabrook E, Kim H, Yoo SSFunctional and Diffusion Tensor Magnetic Resonance Imaging of the Sheep Brain.BMC Vet Res. 2015 Oct 14;11:262. doi: 10.1186/s12917-015-0581-8.

Lévy F, Batailler M, Meurisse M, Keller M, Cornilleau F, Moussu C, … Differential effects of oxytocin on olfactory, hippocampal and hypothalamic neurogenesis in adult sheep. Neuroscience Letters. 2019, 713, 134520

Marino and Merskin, Intelligence, complexity, and individuality in sheep Animal Sentience 2019, 206

McGregor KM, Carpenter H, Kleim E, Sudhyadhom A, White KD, Butler AJ, Kleim J, Crosson B. Motor map reliability and aging: a TMS/fMRI study. Exp Brain Res. 2012 May;219(1):97–106. doi: 10.1007/s00221-012-3070-3.

Mitchell NL, Russell KN, Wellby MP, Wicky HE, Schoderboeck L, Barrell GK, Melzer TR, Gray SJ, Hughes SM, Palmer DN. Longitudinal In Vivo Monitoring of the CNS Demonstrates the Efficacy of Gene Therapy in a Sheep Model of CLN5 Batten Disease. Mol Ther. 2018 Oct 3;26(10):s2366–2378. doi: 10.1016/j.ymthe.2018.07.015.

Peruzzo Denis, Bertoldo Alessandra, Zanderigo Francesca and Cobelli Claudio, Automatic selection of arterial input function on dynamic contrast-enhanced MR images”, Computer methods and programs in biomedicine, 104:e148–e157 (2011).

Pieri V, Trovatelli M, Cadioli M, Zani DD, Brizzola D, Ravasio G, Fabio Acocella F, Di Giancamillo M, Malfassi L, Dolera M, Riva M, Bello L, Falini A, Castellano A. *In vivo* Diffusion Tensor Magnetic Resonance Tractography of the Sheep Brain: An Atlas of the Ovine White Matter Fiber Bundles. Front Vet Sci. 2019 Oct 16;6:345. doi: 10.3389/fvets.2019.00345

Ramadoss J, Tress U, Chen WJ, Cudd TA. Maternal adrenocorticotropin, cortisol, and thyroid hormone responses to all three-trimester equivalent repeated binge alcohol exposure: ovine model Alcohol. 2008 May;42(3):199–205. doi: 10.1016/j.alcohol.2007.12.004

Reimann HM, Niendorf T - The (un) conscious mouse as a model for human brain functions: key principles of anesthesia and their impact on translational neuroimaging Frontiers in Systems Neuroscience, 2020 - frontiersin.org

Restom K, Bangen KJ, Bondi MW, Perthen JE, Liu TT.Cerebral Blood Flow and BOLD Responses to a Memory Encoding Task: A Comparison Between Healthy Young and Elderly Adults. Neuroimage 2007 Aug 15;37(2):430–9. doi: 10.1016/j.neuroimage.2007.05.024.

S Sonnay, J Poirot, N Just, AC Clerc, R Gruetter, G Rainer, JMN Duarte (2018). Astrocytic and neuronal oxidative metabolism are coupled to the rate of glutamate–glutamine cycle in the tree shrew visual cortexGlia 66 (3), 477–491

Schiffner R, Bischoff SJ, Lehmann T, Rakers F, Rupprecht S, Matziolis G, Schubert H, Schwab M, Huber O, Lemke C, Schmidt M. Underlying mechanism of subcortical brain protection during hypoxia and reoxygenation in a sheep model - Influence of α1-adrenergic signalling. PLoS One. 2018 May 29;13(5):e0196363. doi: 10.1371/journal.pone.0196363.

Stypulkowski PH, Stanslaski SR, Denison TJ, Giftakis JE. Chronic evaluation of a clinical system for deep brain stimulation and recording of neural network activity. Stereotact Funct Neurosurg. 2013;91(4):220–32. doi: 10.1159/000345493.

Swallow KM, Braver TS, Snyder AZ, Speer NK, Zacks JM. Reliability of functional localization using fMRI. Neuroimage. 2003;20(3):1561–77. doi: 10.1016/s1053-8119(03)00436-1

van Alst TM, Wachsmuth L, Datunashvili M, Albers F, Just N, Budde T, Faber C Anesthesia differentially modulates neuronal and vascular contributions to the BOLD signal. Neuroimage 2019,195, 89–103

Wahl RU. Could oxytocin administration during labor contribute to autism and related behavioral disorders?-A look at the literature. Med Hypotheses. 2004;63(3):456–60. doi: 10.1016/j.mehy.2004.03.008.

Can Wu, Amir R Honarmand, Susanne Schnell, Ryan Kuhn, Samantha E Schoeneman, Sameer A Ansari, James Carr, Michael Markl, Ali Shaibani Age-Related Changes of Normal Cerebral and Cardiac Blood Flow in Children and Adults Aged 7 Months to 61 Years. J Am Heart Assoc. 2016 Jan 4;5(1):e002657. doi: 10.1161/JAHA.115.002657.

Zanderigo Francesca, and Bertoldo Alessandra and Pillonetto Gianluigi and Cobelli Claudio, Nonlinear stochastic regularization to characterize tissue residue function in bolus-tracking MRI: assessment and comparison with SVD, block-circulant SVD, and Tikhonov, IEEE Transactions on Biomedical Engineering, 56(5):1287--1297 (2009).

